# Mechanical coupling of supracellular stress amplification and tissue fluidization during exit from quiescence

**DOI:** 10.1101/2021.10.19.464930

**Authors:** Emma Lång, Christian Pedersen, Anna Lång, Pernille Blicher, Arne Klungland, Andreas Carlson, Stig Ove Bøe

## Abstract

Cellular quiescence is a state of reversible cell cycle arrest that is associated with tissue dormancy. Timely regulated entry into and exit from quiescence is important for processes such as tissue homeostasis, tissue repair, stem cell maintenance, developmental processes and immunity. Here we show that quiescent human keratinocyte monolayers contain an actinomyosin-based system that facilitates global viscoelastic flow upon serum-stimulated exit from quiescence. Mechanistically, serum exposure causes rapid amplification of pre-existing contractile sites leading to a burst in monolayer stress that subsequently drives monolayer fluidization. The stress magnitude after quiescence exit correlates with quiescence depth, and a critical stress level must be reached to overcome the cell sheet displacement barrier. The study shows that static quiescent cell monolayers are mechanically poised for motility and identifies global stress amplification as a mechanism for tissue fluidization.

## Introduction

Quiescence refers to a state of cell cycle arrest in which cells are retained in a standby mode, ready to re-enter the cell cycle upon activation by a given physiological stimuli. The pool of quiescent cells in the human body is typically represented by tissue-specific stem and progenitor cells, naïve immune cells, fibroblasts and epithelial cells ^1^. In addition, certain cancer cells have the ability to evade cancer therapy by entering a dormant quiescence-like state ^1^. Accordingly, careful regulation of entry into and exit out of quiescence is important for several physiological processes such as tissue homeostasis and repair, stem cell maintenance, immunity, reproduction and development ^1^. Quiescent cells are required to maintain a high level of preparedness in order to facilitate rapid activation of specialized cell functions once cell division is stimulated. In agreement with this, quiescent stem cells and naïve immune cells have been shown to possess multiple epigenetic and post-translation mechanisms that facilitate rapid expression of linage-specific genes following stimulation of quiescence exit ^2–14^. However, little is known about cellular mechanisms that control biomechanical changes in cells as they enter the cell cycle after prolonged periods in quiescence.

Quiescence exit is frequently associated with activation of cell motility. For example, quiescent stem and naïve immune cells migrate out of their niches in response to cell cycle activation in order to support tissue homeostasis, repopulate injured tissue or to perform immune surveillance at distal locations ^15–18^. In addition, reawakening of dormant quiescent cancer cells can cause tumor relapse and formation of metastases years after remission ^19^. In multilayered epithelial tissue, like the skin, exit from quiescence during homeostasis is associated with lateral migration to suprabasal regions, while skin injury evokes massive reawakening of basally localized keratinocytes concomitant with activation of cell sheet displacement by collective migration to restore damaged epidermal surfaces ^20–23^. The strong correlation between quiescence exit and cell migration in multiple physiological settings suggest the existence of mechanisms that link quiescence exit to activation of cell motility.

In this study we have investigated a mechanical link between quiescence exit and cell migration using the immortalized human keratinocyte cell line HaCaT. This cell line has been shown to support long-range cell sheet displacements through collective cell migration following serum-activated cell cycle re-entry ^24^. Using traction force microscopy we found that quiescent monolayers generate cores of weak contractility that are amplified shortly after serum stimulation to create a coordinated global burst of traction forces and stress throughout the cell monolayer. These forces are actinomyosin-dependent and mechanically coupled to large-scale coordinated cell sheet displacements through flow as a liquid-like matter. A theoretical model for the viscoelastic tissue shows that force amplification within randomly distributed contractility sites is sufficient to direct coordinated cell motion. Finally, the magnitude of mechanical forces created during quiescence exit correlates with quiescence depth and the extent of cell sheet displacements. Our study provides evidence that quiescent keratinocyte monolayers possess mechanical preparedness for motility and suggest that stress amplification is a generic mechanism for tissue fluidization.

## Results

### Quiescence-dependent cell sheet displacements emerge through collective migration and large-scale deformations

In a previous study quiescent HaCaT keratinocytes were observed to transform from static to motile cell sheets following exposure to serum. The serum-activated motions in these experiments are coordinated over length scales exceeding several millimeters and depend on prior activation of quiescence through serum depletion ^24^. In the present study we imaged mCherry-tagged nuclei within HaCaT keratinocyte monolayers seeded in 96-well plates using an automated high-content imaging system. Using this approach, we were able to analyze cell sheet velocity, cell density and cell sheet thickness based on multiple cell monolayer samples confined to a defined circular surface (7.0 mm in diameter) (Fig. 1a). We observed limited cell motion within confluent cell monolayers that had been plated in serum-containing medium 20 hours (h) prior to imaging and cell monolayers that had been subjected to an additional two days of serum depletion prior to imaging (Fig. 1b, Supplementary Video 1). This is consistent with previous reports showing that epithelial cells undergo jamming and immobilization at high densities when confined to a fixed area ^25,26^. In contrast, serum re-exposure of serum-depleted monolayers led to quiescent-dependent cell sheet displacements (Fig. 1b, Supplementary Video 1).

**Figure 1:**
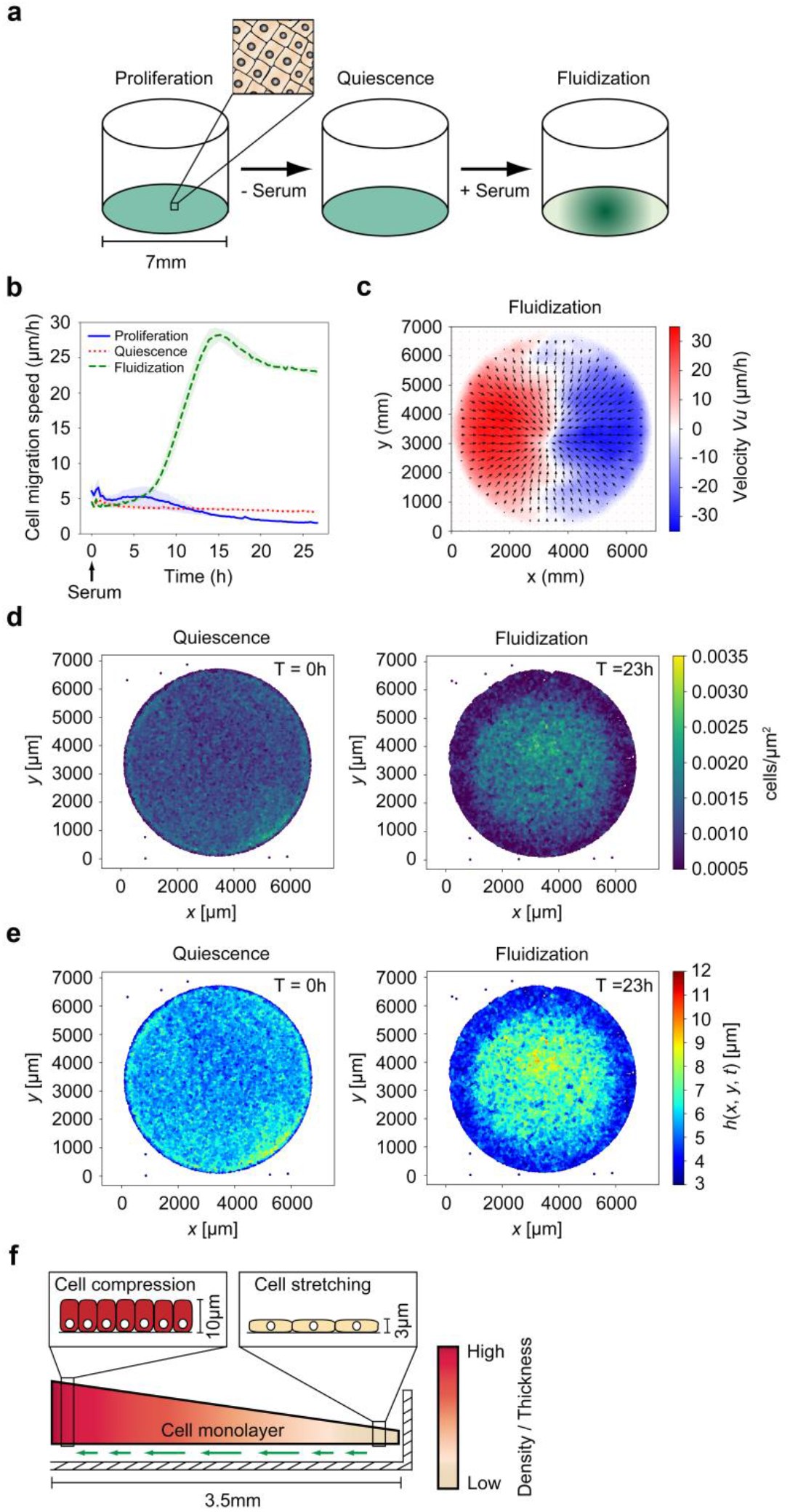
Quiescence-dependent cell sheet displacements by collective migration and large-scale deformation. **a**, Schematics of the methodology used for generation of quiescence-dependent cell sheet displacement. **b**, Cell migration speed in confluent monolayers without treatment (proliferation), monolayers subjected to serum depletion for 48 h (quiescence), and monolayers subjected to serum depletion for 48 h followed by serum re-stimulation (fluidization). Data are based on PIV analysis and show average migration speed across the monolayer surface ± standard deviation (SD), n = 8 separate monolayers per sample. See also Supplementary Video 1. **c**, PIV-derived vector fields 18 h after serum re-stimulation. Average values of 8 separate monolayers are shown. **d**, Cell density at 0 h and 23 h after serum re-stimulation. See also Supplementary Video 2. **e**, Cell sheet thickness at 0 h and 23 h after serum re-stimulation. See also Supplementary Video 2. **f**, Schematic showing cell sheet displacement (green arrows) and deformation through viscoelastic flow. Cross section represents one half (3.5 mm) of the confluent cell sheet in a well of a 96-well plate.

The serum-activated quiescent monolayers exhibited a remarkably uniform cell migration pattern within the 96-well plate that was characterized by collective flow of material away from the monolayer edges towards the central region of the well (Fig. 1c-e, Supplementary Video 2). Analysis of cell density revealed a progressive increase in cell density at the center concomitant with decreasing cell density near the monolayer edges (Fig. 1d, Supplementary Video 2). To analyze monolayer compression and extension we developed an algorithm for mapping the monolayer thickness based on its cellular density (Supplementary Fig. 1). Mapping of monolayer thickness in serum activated quiescent cell sheets, revealed a progressive increase in thickness at monolayer centers (maximum cell sheet height of 10 μm) and a compensatory decrease in monolayer thickness at the cell sheet edges (minimal cell sheet height of 3 μm) following serum-induced exit from quiescence (Fig. 1e, Supplementary Video 2). We also investigated a potential remodeling of actin at the basal cell surface using live confocal microscopy of LifeAct-expressing HaCaT cells. We observed a rapid disappearance of stationary actin-enriched foci at the basal side during the first 3 h after serum re-stimulation (Supplementary Video 3). This was followed by the appearance of dynamic actin-enriched “feet” that coincides with activation of cell sheet displacement 5 to 8 h after serum stimulation (Supplementary Video 4). Combined, these results show that serum-activated quiescent cell sheets mediate large-scale displacements through collective cell migration and redistribution of mass, dynamic features that are consistent with viscoelastic flow (Fig. 1f).

### Serum activation of quiescent cells generates an immediate burst of traction forces and intercellular stress through amplification of pre-existing contractile sites

To investigate the mechanical forces that regulate quiescence-dependent collective cell migration, we extracted traction forces and the intercellular stress in cell monolayers using traction force microscopy (TFM). To achieve this we plated HaCaT keratinocytes on a soft (4kPa) substrate consisting of collagen IV-coated polyacrylamide containing fluorescently labeled beads as fiducial markers. We analyzed multiple microscopy fields (1.3×1.3 mm each) within 12-well glass bottom plates. Cell sheets plated on soft acrylamide substrates exhibited a quiescence-dependent migration phenotype similar to that observed for cells grown on a glass surface. We noted a lower maximal migration speed for cells plated on acrylamide (15-20 μm/h) compared to cells plated on glass (25-35 μm/h), a difference that could be attributed to differences in substrate softness. Analysis of cell-mediated substrate displacements at time points before and after serum stimulation revealed an instant increase in traction forces and intercellular stresses across the entire confluent cell sheet after serum stimulation (Fig. 2a and b, Supplementary Video 5). This burst in mechanical forces peaked between 10 and 30 minutes (min) after serum exposure and progressively decayed for a period of more than 20 h concomitant with activation of cell sheet motility (Fig. 2c-g). The inverse correlation between monolayer tension and cell sheet displacement activity after an initial burst in traction force magnitude was unexpected since most studies performed on single cells and cell collectives suggest a positive correlation between traction forces and cell motility ^27–29^.

**Figure 2:**
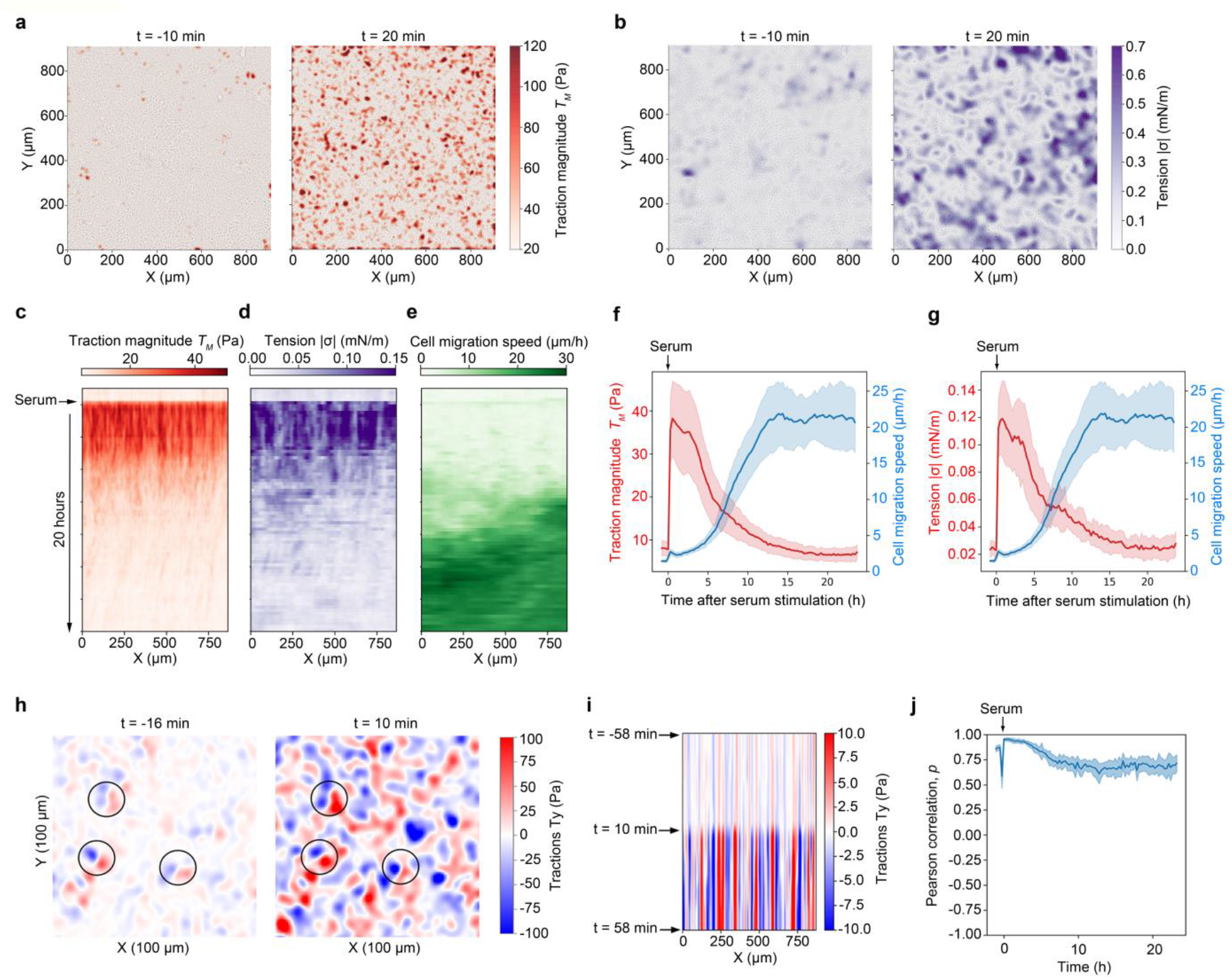
Serum activation of quiescent monolayers is accompanied by activation of global stress through rapid amplification of pre-existing stress centers. **a**, Traction force maps generated 10 min before (left panel) and 20 min after (right panel) serum activation of serum-depleted monolayers. See also Supplementary Video 3. **b**, Tension maps 10 min before (left panel) and 20 min after serum activation of serum-depleted monolayers. See also Supplementary Video 3. **c**-**e**, Kymographs showing traction magnitude (**c**), intercellular tension (**d**), and migration speed (**e**) before and after serum stimulation of a confluent quiescent cell sheet. A representative microscopic field 48 min before and 20 h after serum stimulation is shown. **f**,**g**, Traction magnitude (**f**) and intercellular tension (**g**) relative to cell migration speed over time. The plotted lines show average ± SD, n=6 separate microscopic fields. **h-j**, Pre-existing local tractions in quiescent cell sheets are amplified by serum re-stimulation. **h**, Map showing *T_y_* traction force component 16 min before and 10 min after serum re-stimulation. Examples of amplified traction sites are highlighted by circles. **i**, Kymograph displaying the traction force *T_y_* component before and after serum re-stimulation of a confluent quiescent cell sheet. **j**, Plot showing time evolution of the Pearson correlation coefficient. Each of the plotted values represents the Pearson correlation between adjacent frames. n=6 separate microscopic fields.

We next analyzed the relative positions of traction force centers before and after serum stimulation. We found that local serum-stimulated tractions frequently arise from weaker tractions already present in the quiescent unstimulated cell sheet (Fig. 2h and i). In agreement with this, we observed a positive Pearson correlation coefficient between traction force maps generated before and after serum stimulation (Fig. 2j). This result shows that quiescent monolayers are poised for a high global stress by means of numerous weak local tractions that can be amplified upon exposure to serum-born growth factors.

We also analyzed the individual traction force components in x-direction (*T_x_*) and in y-direction (*T_y_*) within microscopic fields with the aim of identifying potential biases in traction force orientation. Inspection of the traction components did not reveal a preferential direction of force orientation across larger microscopy areas (Supplementary Fig. 2a and b). In addition, plotting the average of the traction force component *T_x_* against increasing microscopic field size suggested that the average traction forces in the x or y direction quickly approached zero for microscopic fields larger than 100×100 μm (Supplementary Fig. 2c). Thus, the orientation of local traction forces before and after serum stimulation is largely random relative to the direction of subsequent cell sheet displacement.

### Stress amplification is actinomyosin dependent and coupled to cell sheet displacement

Actinomyosin is known to regulate tractions and stress in cells during single cell or collective cell migration. To investigate the role of actinomyosin in quiescence-dependent collective migration we serum-activated quiescent cell sheets grown on soft acrylamide substrates in the presence or absence of the Rho-associated kinase (ROCK) inhibitor Y-27632 or the non-muscle myosin inhibitor Blebbistatin. The presence of these inhibitors led to a significant reduction in traction force amplification and cell sheet velocity after serum activation (Fig. 3a and b).

**Figure 3:**
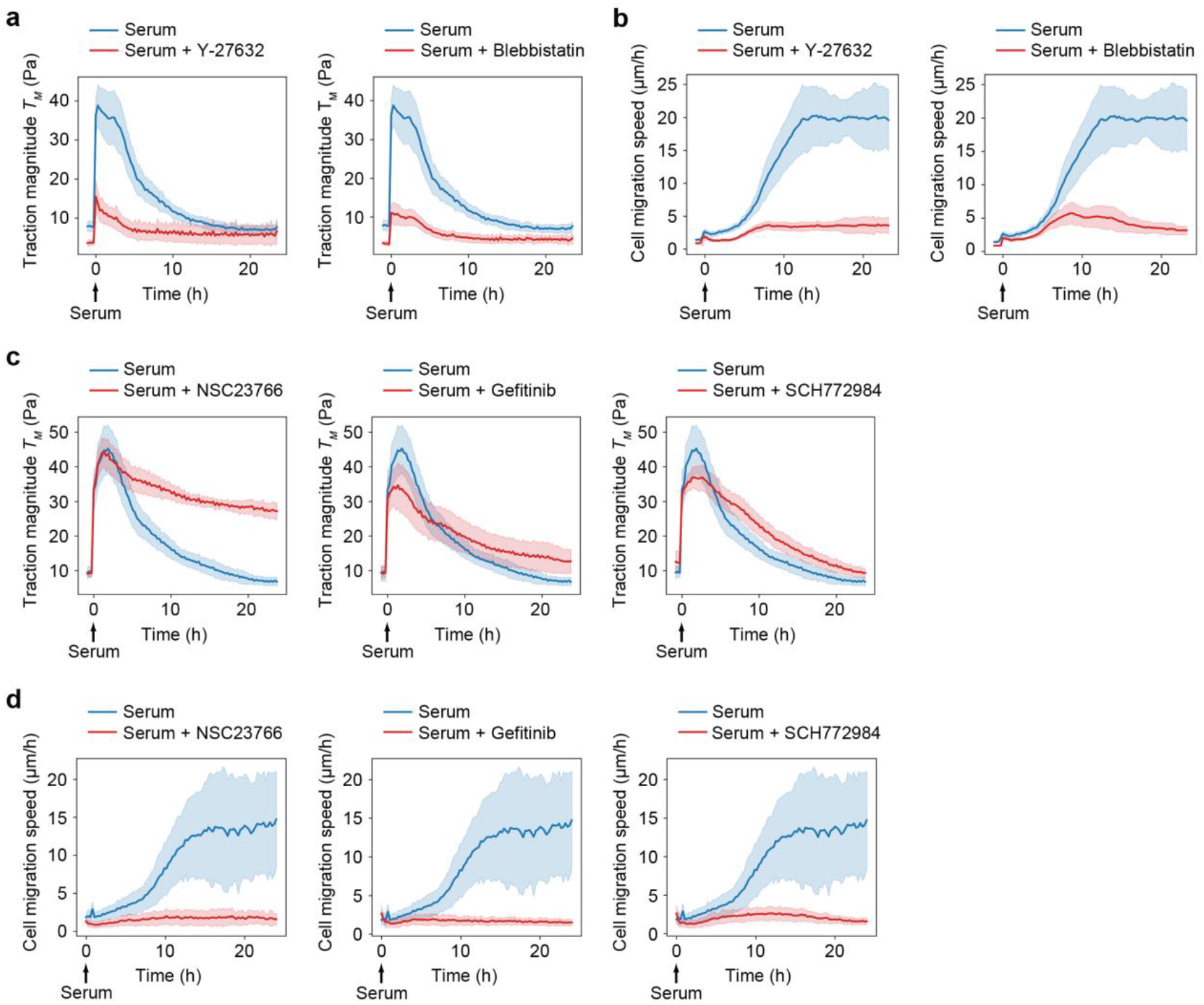
Dependency between global stress amplification and cell sheet displacement. **a**, Traction force magnitudes during serum activation in the presence of Y-27632 or Blebbistatin. **b**, Cell migration speed during serum activation in the presence of Y-27632 or Blebbistatin. **c**, Traction force magnitudes generated in the presence of collective cell migration inhibitors for Rac1, EGFR, and ERK. **d**, Cell migration speed generated in the presence of collective cell migration inhibitors for Rac1, EGFR, and ERK. **a**-**d**, All graphs show mean values ± SD from 8 separate microscopy fields.

We next examined a potential link between the amplified mechanical forces and cell migration by performing pharmacological inhibition of Rac1, EGFR and ERK, protein components that play critical roles in mediating collective cell migration in epithelial monolayers ^24,30^. Inhibitors against these proteins caused a reduction in traction force decay rates after the initial serum-mediated traction force amplification (Fig. 3c). For cell cultures treated with the Rac1 inhibitor the traction force amplification after serum stimulation seemed unaffected, while the subsequent traction force decay was strongly reduced compared to that of control-treated cells (Fig. 3c). For cells treated with EGFR or ERK inhibitors the effect on traction force decay was significant but less pronounced compared to Rac1-treated cells, possibly due to a dual role of these inhibitors in interfering with both traction force amplification and cell migration (Fig. 3c). We confirmed that all three inhibitors prevented quiescence-induced collective cell migration on the soft acrylamide substrate (Fig. 3d). These observations support the notion that the decay in tissue stress is coupled to activation of collective cell migration. Combined, these results suggest that exposure of quiescent monolayers to serum stimulates actinomyosin-mediated monolayer tension and that the subsequent large-scale cell sheet displacements are driven by monolayer tension decay.

### Theory predicts serum-induced stress amplification as a key component for global cell sheet displacement

To understand how the stress distribution in the cell sheet can lead to collective migration of the cells, we develop a minimal mathematical model based on an active viscoelastic gel theory ^31–37^ (see methods and supporting information (SI)). The model encompass the most essential physical components in the cell layer i.e. elastic deformations, cell-substrate friction and viscous friction, in addition to an active stress induced by the actinomyosin (described by the field c(x, t)). The experimental conditions are mimicked in the numerical simulations when solving the mathematical model: we use a circular domain, with an initial random distribution of the actinomyosin giving the pre-stressed state of the sheet. When there is a low level of local contractility in the sheet, we observe little net motion and a global cell sheet displacement fail to appear (Fig. 4a-c, Supplementary Video 6). We then tested how the increased level of pre-stress affects the cell sheet dynamics by systematically increasing the initial magnitude of c(**x**, t=0). We found that a progressive increase in monolayer pre-stress correlates with increased activation of radial displacement from the edges of the monolayer towards the center (Fig. 4d-f; Supplementary Video 7). Thus, by using a sufficiently high initial pre-stress magnitude, the model recapitulates the cell sheet dynamics observed following exposure of quiescent cell monolayers to serum.

**Figure 4:**
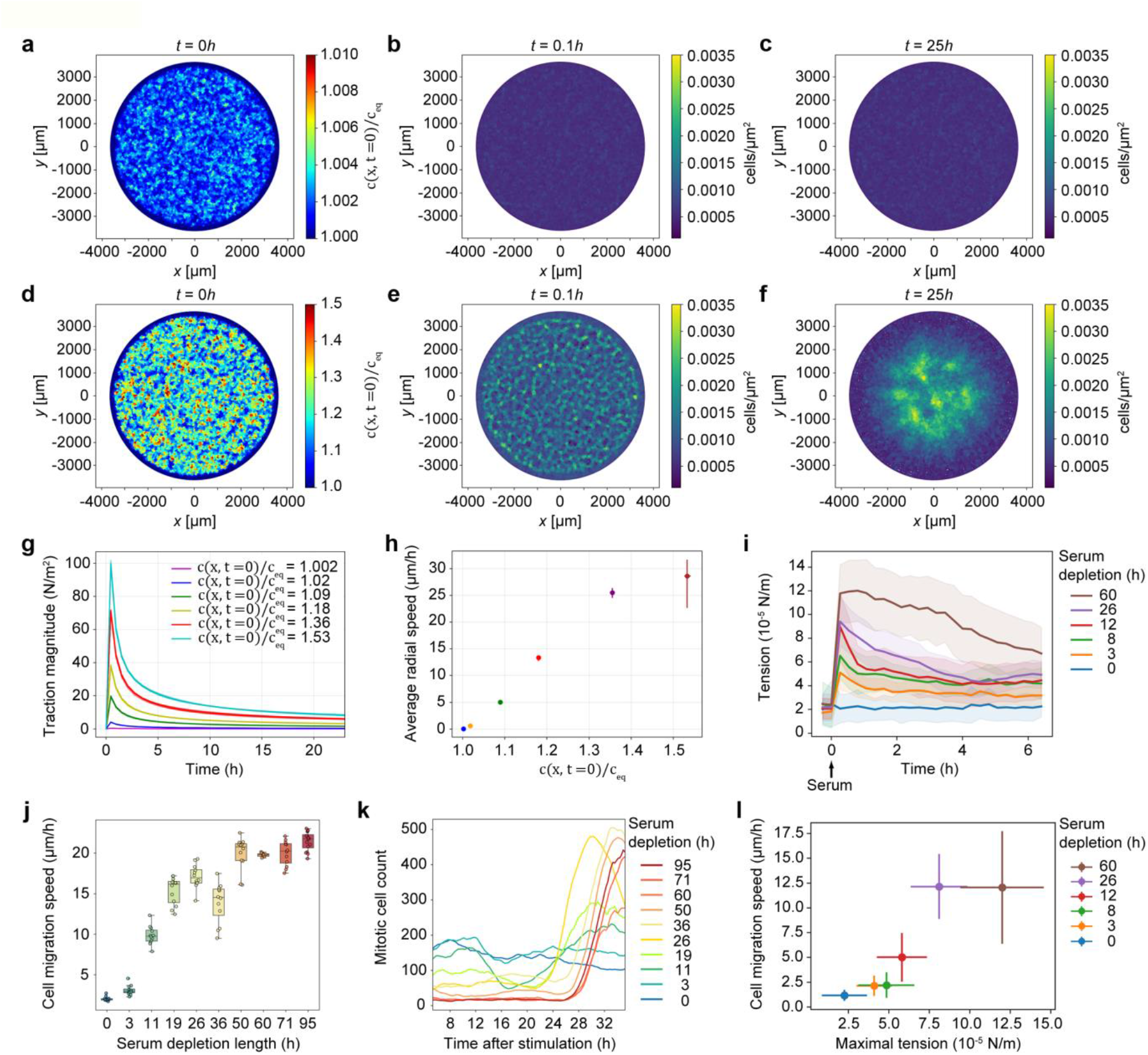
Theory predicts serum-induced stress amplification as a key component for global cell sheet displacement. **a**, Normalized initial concentration c(x, t=0)/c_eq_ with a pre-factor 0.01 is used to calculate the results shown in panels **b**-**c**. See also Supplementary Video 4. **b**, Monolayer displacement field u(x, t) at time t=0.1h. Small local contractions are formed at points that have a high initial concentration. **c**, Monolayer displacement field u(x, t) at time t=25h. At long times the monolayer remains in a homogenous state and we observe no collective migration to form a central contraction centre. **d**, Normalized initial concentration c(x, t=0)/c_eq_ with a pre-factor 1.0 used to calculate the results shown in panels **e**-**f**. See also Supplementary Video 5. **e**, Monolayer displacement field u(x, t) at time t=0.1h. Strong local contraction centres are formed at points that have a high initial concentration. **f**, Monolayer displacement field u(x, t) at time t=25h. At long times we observe a collective migration towards the centre of the monolayer, forming one large central contraction centre. **g**, Traction magnitude as a function of time. **h**, Average radial speed in the cellular monolayer as a function of the normalized initial concentration c(x, t=0). The parameters used in the numerical simulations (a-h); E = 4000 Pa, h_eq_ = 8 μm, D = 10^−9^ m^2^ s^−1^, Γ= 100 Ns m^−3^, α = 12 Pa, ***β*** = 0.24 s^−1^, *τ_c_* = 13000 s, are further described in the SI. **i**, Time evolution of intercellular tension produced after different time periods of serum depletion. Graph shows mean values ± SD, n=10-25 separate microscopic fields from 2 independent experiments. **j**, The effect of serum deprivation length on cell migration. Monolayers in 96-well plates were subjected to serum deprivation for the time periods indicated and subsequently re-stimulated with serum. Migration speed is plotted as the mean velocity between 8 and 50 h after serum re-stimulation. n=8-16 separate microscopic fields from 2 independent experiments. **k**, Estimation of the cell division frequency after different length of serum depletion prior to serum re-stimulation using automated detection of mitotic cells. Graph shows moving average of mitotic cell counts and is representative of two independent experiments. **l**, Plot showing cell sheet displacement magnitude versus intercellular tension. Maximal tension is represented by the tension measured at 1 h after serum re-stimulation. Average speed is represented by the average speed of all time points between 8 and 20 h after serum re-stimulation. The plot uses the same set of raw data as for panel **i**.

To further quantify the influence of the pre-stress, we extract the averaged traction force (Fig. 4g) and the averaged radial velocity (Fig. 4h) for the different initial magnitudes of c(**x**, t=0). We observed a steep increase in the traction force magnitude that decays with time (Fig. 4g). Furthermore, the radial velocity appears to scale linearly with the magnitude of initial stress until a threshold value of the active actinomyosin is approached whereby the velocity stagnates (Fig. 4h). A linear response in radial velocity with respect to the magnitude of c(**x**, t=0) and the length over which c(**x**, t=0) varies is predicted from scaling analysis of the mathematical model (see SI, Supplementary Fig. 3). However, as the cellular density at the center of the monolayer increases the elastic compression will balance the concentration gradient and thus lead to a stagnation and subsequent reduction in both c(x, *t*) and in the cells averaged radial velocity. Combined, the theoretical predictions point to the level of cell sheet pre-stress as the key feature for activation of global monolayer displacements.

To verify the effect of different stress levels on cell sheet displacement in experiments, we reasoned that the monolayer stress can be tuned by adjusting the length of the serum deprivation period prior to serum re-exposure. We therefore examined the relative relationship between the serum depletion length and the level of intercellular tension amplified after serum stimulation. We found that the serum depletion length correlates with the level of intercellular stress produced upon serum re-stimulation (Fig. 4i). Serum depletion length was also found to correlate with cell velocity after serum re-stimulation (Fig. 4j, Supplementary Fig. 4). To analyze the effect of serum depletion length on quiescence depth we monitored the surge of cell divisions that occur between 24 and 30 h after re-exposure of quiescence cells to serum. We found that the serum-deprivation length correlates with the time required for cells to enter mitosis after serum exposure (Fig. 4k). This suggests that the level of monolayer stress after serum re-stimulation correlates with quiescence depth. Finally, by plotting tension values measured in serum-stimulated monolayers subjected to different serum depletion lengths against cell sheet velocities, we found that a threshold value of tension between 0.05 and 0.12 mN/m is required in order to achieve effective collective migration (Fig. 4l). Together, predictions from a minimal theoretical model and experimental observations suggest that the transition of quiescent monolayers from rigid immobile to dynamic viscoelastic cell sheets is mediated trough rapid and coordinated amplification of numerous traction force centers dispersed throughout the monolayer. Indeed, both the theoretical model as well as the experiments shows that a monolayer has to overcome a critical level of global stress in order to achieve effective viscoelastic flow.

### Conclusions and outlook

Many cellular processes, including tissue development and repair, require both cell cycle re-entry and tissue fluidization. In agreement with this, the present study identifies a mechanism that couples cell sheet fluidization to exit from quiescence. Mechanistically, the quiescence-dependent activation of viscoelastic flow occurs through global amplification of monolayer stress that subsequently drives cell migration. The requirement of a critical stress level for activation of tissue fluidization is consistent with previous studies showing that motion in developing tissues is driven by supracellular stress alterations^38,39^. Thus, tissue stress amplification may represent a generic mechanism for transformation of tissues from a solid to a liquid-like state.

The observation that quiescent monolayers contain cores of pre-installed weak contractility that can undergo force amplification after serum exposure suggest that dormant tissues are mechanically prepared for fluidization. Such tissue preparedness may have important consequences for tissue regeneration. For example, defects in tissue-scale stress amplification during quiescence exit shortly after injury may lead to impaired healing. Further studies should clarify if quiescence-dependent fluidization is linked to the pathology of non-healing wounds.

## Methods

### Cells

The immortalized human keratinocyte cell line HaCaT ^40^ and its derivatives were grown in Iscove’s modified Dulbecco’s medium (IMDM, MedProbe) supplemented with 10% fetal bovine serum (FBS, Thermo Fisher Scientific) and 90 U/ml penicillin/streptomycin (PenStrep, Lonza). For serum depletion and serum re-activation we used the same medium, but with 0 and 15% FBS, respectively. HaCaT cells expressing mCherry-tagged Histone H2B has been described previously ^24^. HaCaT cells stably expressing LifeAct-RFP were produced by using recombinant lentivirus particles obtained from Ipidi.

### Drug treatment

Inhibitors used in the study: Y-27632 (Y0503, Merck; used at 5μg/ml final concentration), Blebbistatin (B0560, Merck; used at 2 μg/ml final concentration), Gefitinib (Y0001813, Merck; used at 5 μM final concentration), SCH772984 (S7101, Selleck Chemicals; used at 10 μM final concentration), NSC23766 (SML0952, Merck; used at 100μM final concentration). Cells that had undergone serum depletion for 48 h were pretreated with drugs for 1 h in serum-depleted medium before they were placed under the microscope. Images of cells and polyacrylamide-embedded beads were then acquired at 3 time points (intervals of 16 min) before serum activation by adding 1 volume of medium containing 30% serum and 2x concentrations of respective drugs. Control cells were treated the same, but without drugs.

### High content imaging of monolayers confined to the bottom surface of 96-well plates

HaCaT mCherry-Histone H2B cells were seeded in 96-well glass bottom plates (Greiner Sensoplate (M4187-16EA, Merck)) at 75 000 cells per well. Cells were consistently seeded 18 to 22 h prior to serum depletion or image acquisition, also when different lengths of serum depletion were compared. Live cell imaging of monolayers was performed using the ImageXpress Micro Confocal High-Content Microscope controlled by the MetaXpress 6 software (Molecular Devices). Image acquisition was carried out in widefield mode using a 4x 0.2 NA PLAPO air objective at pixel binning 2, filter set for detection of mCherry fluorescence, and an environmental control gasket that maintain 37°C and 5% CO_2_. Four tiled images per well (covering the entire well surface) were acquired for a total time period of 30 to 50 h using a time interval of 16 min between frames. Acquired time lapse movies were then processed by tiling (using the MetaXpress software) to generate movies covering the entire well bottom surface.

### Preparation of PAA gel substrates

Collagen IV-coated polyacrylamide (PAA) gels were prepared in 12-well glass bottom multiwell plates (P12G-1.5-14-F, MatTek Corporation) using a protocol modified from Serra-Picamal *et al*.^41^. The glass surfaces were activated by adding Bind-silane (GE17-1330-01, Merck):MQ-H_2_O (Milli-Q^®^ Reference Water Purification System): acetic acid at a 1:12:1 ratio for 4 min followed by two quick washes in 1x phosphate buffered saline (PBS). 4.0 kPa polyacrylamide gels were prepared by mixing 93.75 μl 40% (7.5%) acrylamide solution (1610140, Bio-Rad), 37.5 μl 2% Bis-solution (161-0142, Bio-Rad), 2 μl (0.4%) Carboxylate-modified fluorescent beads (FluoSpheres^™^ Carboxylate-Modified, 0.2 μm, 580/605; F8810, Thermo Fisher Scientific), 2.5 μl Ammonium Persulfate (0.05%, Bio-Rad), 0.25 μl TEMED (0.05%,161-0801, Bio-Rad) and 364 μl MQ-H_2_O. 10 μl freshly prepared gel solution was placed as droplets at the bottom of each glass bottom well and subsequently overlaid by GelBond film (80112933, Cytiva) cut to a circle, 12 mm in diameter, hydrophobic side facing down.

The 12-well plate was placed in an inverted position for 40 min while the gel polymerized. Following gel polymerization, 2 ml 10x PBS was added to each well for 40 min prior to removal of the GelBond support. Gels were then washed 2x 4 min in 1x PBS and subsequently treated with 40 μl Sulfo-SANPAH (sulfosuccinimidyl 6-(4’-azido-2’-nitrophenylamino) hexanoate (22589, Thermo Fischer Scientific) for 4 min under UV light. The UV-treated Sulfo-SANPAH was then replaced with fresh Sulfo-SANPAH, and treated with UV light for another 6 min. Following Sulfo-SANPAH cross linking, cells were washed 2x 4 min in PBS and subsequently overlaid with 0.1 mg/ml collagen IV (C7521, Merck) and placed at 4°C overnight (ON). The next day, gels were washed twice in 1x PBS and then stored at 4°C in 1x PBS for a maximum of 2 days before use in Traction force microscopy (TFM) experiments.

### Traction force microscopy (TFM)

The microscope used for TFM was a Zeiss AxioObserver.Z1 equipped with a CO_2_ incubation chamber, a Colibri 7 LED light source, automated stage controlled by the Zen software and a 10x 0.5 NA FLUAR air objective. Cells were seeded on collagen-coated 4.0 kPa PAA gels in 12-well plates at 600 000 cells per well. Following growth for 12 to 20 h in normal growth medium at confluent densities cells were placed on the microscope stage and adopted to the microscope environment for 1 h. Subsequently, 3-4 consecutive frames (16 min intervals between frames) were captured in order to acquire a “before stimulation” reference. Imaging was then paused while a final concentration of 15% FBS was added in order to serum-activate the cell sheets. For each imaging time point a total of 4 sites per well was captured. The 10x objective generated a field of view of 1331×1331 μm and for each site a stack of 6 z-planes was collected with a z-step size of 2.2 μm. For each z-stack, the middle z-plane was focused at the center of the gel surface. The time interval between frames was 8 min and two channels were captured, one for the beads (580/605 nm) and one for transmitted light to capture cell movements. Intervals between frames were set to 16 min. At the end of each image acquisition series cells were subjected to trypsin treatment for 1 h in order to capture a final reference frame of unstrained fluorescent beads.

To calculate traction forces each of the z-stacks from individual sites were projected using a maximal intensity projection (MIP) algorithm. Subsequently, all images in a time series were subjected to image registration using the descriptor-based series registration plugin in Fiji ImageJ ^42^. Bead displacement traction forces and tension maps were calculated using open source codes available from pyTFM ^43^.

### Kymographs

Spatiotemporal diagrams (kymographs) were generated based on time lapse images generated by particle image velocimetry (PIV) or TFM using an *in-house* Python-based script. One-dimensional images were generated by averaging pixel values along the Y-axis. Two dimensional kymographs were then created by stacking the one dimensional images.

### Monolayer thickness measurements

HaCaT LifeAct-mCherry/EGFP-Histone H2B cells were seeded in 6 cm MatTek dishes at confluent densities. Volumetric stacks of living cell monolayers were generated at various cell densities following serum depletion for 48 h and subsequent serum re-activation for 18 h. The microscope used was a Leica SP8 confocal microscope equipped with a 40x oil immersion objective, an incubation chamber maintaining 37°C and 5% CO_2_, Hybrid (HyD) detectors and the LAS X software. For each field of view (291×291 μm) the average monolayer thickness was calculated by measuring the depth of the mCherry signal at four randomly selected positions, while cell density in the same image was estimated based on EGFP-labeled Histone H2B using the Find Maxima function in ImageJ. By fitting a linear polynomial to the average thickness data we obtained a function that describes the correlation between monolayer thickness and cell sheet density. This function was used to calculate the cell sheet thickness in high-content imaging experiments with 96-well plates, based on monolayer density measurements acquired with the Find Maxima function.

### Visualization of basal actin dynamics

HaCaT RFP-LifeAct cells were seeded at confluent cell densities in 35 mm bottom glass dishes from MatTec. Cells were subjected to 48 h of serum depletion followed by serum re-stimulation. Imaging of the basal cell surface after stimulation was carried out using a Leica SP8 confocal microscope equipped with a 40x oil immersion objective, an incubation chamber maintaining 37°C and 5% CO_2_. Time lapse series of a single Z-position representing the basal cell surface were acquired at a frame rate of 30 seconds between frames.

### Cell division analysis

HaCaT mCherry-Histone H2B cells were seeded in 12-well glass bottom multiwell plates from MatTek (P12G-1.5-14-F, MatTek Corporation) at 800 000 cells per well. Notably, all cells were seeded 12 to 20 h before start of serum depletion. Cells were subjected to different time periods of starvation, followed by serum re-stimulation and wide-field microscopy. Image acquisition was carried out on a Zeiss AxioObserver.Z1 microscope using a 10x 0.5 NA FLUAR air objective, filter set for detection of mCherry fluorescence, and an environmental control gasket that maintain 37°C and 5% CO_2_. Four randomly selected sites per well were acquired for a total time period of 50 h, using a time interval of 16 min between frames. First, mitotic cells in a set of images were annotated and used as input data for development of a deep learning model using the StarDist (2D) network, available in the CoLab notebook ^44^. Second, the deep learning model was further used to detect cell division frequencies in each frame of complete datasets, by running the model in the StarDist plugin of Fiji ImageJ ^45^. In addition, the versatile (fluorescent nuclei) StarDist model, available in Fiji ImageJ ^46^, was used to detect total number of cells in each frame acquired. The results were finally displayed using Python 3.7 scripts.

### Mathematical model

A minimal mathematical model for the epithelial layer, considered to behave as a continuum, is based on an active gel theory ^31–37^ with contributions from elastic deformations, active actinomyosin contractions, viscous friction and cell-substrate friction. The monolayers elastic displacement field is **u**(**x**, t) with **x** the spatial coordinate vector and *t* time. The active actinomyosin is labeled by a concentration field c(**x**, *t*)^47–49^. During cell migration the traction force on the underlying substrate becomes (1) T = Γ∂_t_**u**(**x**, t), with Γ[Ns m^−3^] as the cell-substrate friction coefficient. The monolayer stress consists of a passive elastic stress σ_el_(**x**, t), which is modelled as a linearly elastic material ^50^, and an active stress σ_*α*_(**x**, *t*). We assume a linear relation between c(**x**, *t*) and the active stress ^34^. The dynamics of the monolayer is then given by the force balance, (2) Γ*∂_t_**u**(**x**,t) = h_eq_*E*∇^2^**u**(**x**,*t*) + *h_eq_α*∇c(**x**,t),*where *E* [Pa] is the Young’s modulus, *h_eq_* [m] is the average monolayer thickness and α is the contractile strength. c(**x**, *t*) is described by a convection-diffusion-reaction equation, (3) 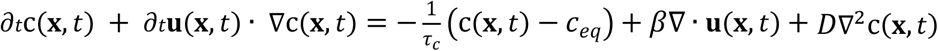, where *τ_c_* [s] is the relaxation timescale towards the equilibrium concentration *c_eq_* [−] that is defined as the concentration at which there are no intracellular contractions, *β*[t^−1^] is the activation rate due to stretching or compression in the monolayer and *D* [m^2^s^−1^] is a diffusion coefficient. A further discussion of the model, the choice of the material parameters and their sensitivity is discussed in the SI.

### Numerical simulations

The mathematical model represented by equations (2)-(3), is solved using a Newton solver from the finite element library FeniCS, where the spatial gradients are discretized by a piecewise linear function. The 2D circular mesh is generated with the software Gmsh using a mesh spacing of 0.015. The boundary conditions used in the simulations are: no displacement **u**(**|x|**=1, t)=**0** and a no flux of c(**x**, t), ∇c(**x**, t)●**n**=0. We seed 10 meshes with an initial condition for c(**x**, t=0) where each spatial mesh point have a 60% chance of being seeded with a random number between 0-1 drawn from a Gaussian distribution. For each initial seeding we simulate the monolayer dynamics by multiplying the concentration with a pre-factor, here [0.01, 0.1, 0.5, 1.0, 2.0, 3.0], before solving (2)-(3). We use an initial non-dimensional time step of 0.00005 which is increased by 5% for each time step until a maximum time step of 0.001 is reached which is then kept constant until a non-dimensional time 1 at which the simulations are terminated. This is sufficiently long for all simulations that display the formation of one large central contraction center has reached its maximum contraction. When the simulations are allowed to further continue in time the layer eventually flattens with a uniform c(**x**, t)=c_eq_ field.

### Averaged radial velocity

The radial velocity in the monolayer is obtained by calculating the radial displacement at each spatial point from the Cartesian displacement field u(x, t). The average velocity is then calculated by averaging over the 10 simulations for each pre-factor to the initial concentration seeding with the SD being the maximum and minimum values obtained in the 10. Initially, the radial displacement field is random in the sense that it has approximately equal amounts of cells moving towards and away from the centre due to the random concentration seeding and the subsequent formation of small local contraction centres. Therefore, we define the averaged radial velocity to be the averaged radial velocity values obtained in the time starting when less than 1% of the cells are moving away from the centre to the time when the velocity is reduced to 20% of its initial magnitude.

## Acknowledgment

The work was supported by the Research Council of Norway and South Eastern Norway Regional Health Authority.

## References

1 Marescal, O. & Cheeseman, I. M. Cellular Mechanisms and Regulation of Quiescence. Developmental cell 55, 259–271, doi:10.1016/j.devcel.2020.09.029 (2020).

2 Bernstein, B. E. et al. A bivalent chromatin structure marks key developmental genes in embryonic stem cells. Cell 125, 315–326 (2006).

3 Carrelha, J. et al. Hierarchically related lineage-restricted fates of multipotent haematopoietic stem cells. Nature 554, 106–111 (2018).

4 Cheung, T. H. et al. Maintenance of muscle stem-cell quiescence by microRNA-489. Nature 482, 524–528 (2012).

5 Crist, C. G., Montarras, D. & Buckingham, M. Muscle satellite cells are primed for myogenesis but maintain quiescence with sequestration of Myf5 mRNA targeted by microRNA-31 in mRNP granules. Cell stem cell 11, 118–126 (2012).

6 Cui, K. et al. Chromatin signatures in multipotent human hematopoietic stem cells indicate the fate of bivalent genes during differentiation. Cell stem cell 4, 80–93 (2009).

7 Grover, A. et al. Single-cell RNA sequencing reveals molecular and functional platelet bias of aged haematopoietic stem cells. Nature communications 7, 1–12 (2016).

8 Hausburg, M. A. et al. Post-transcriptional regulation of satellite cell quiescence by TTP-mediated mRNA decay. Elife 4, e03390 (2015).

9 Lechman, E. R. et al. Attenuation of miR-126 activity expands HSC in vivo without exhaustion. Cell stem cell 11, 799–811 (2012).

10 Lee, J.et al. Signalling couples hair follicle stem cell quiescence with reduced histone H3 K4/K9/K27me3 for proper tissue homeostasis. Nature communications 7, 1–15 (2016).

11 Liu, L. et al. Chromatin modifications as determinants of muscle stem cell quiescence and chronological aging. Cell reports 4, 189–204 (2013).

12 Llorens-Bobadilla, E. et al. Single-cell transcriptomics reveals a population of dormant neural stem cells that become activated upon brain injury. Cell stem cell 17, 329–340 (2015).

13 van Velthoven, C. T. & Rando, T. A. Stem cell quiescence: dynamism, restraint, and cellular idling. Cell stem cell 24, 213–225 (2019).

14 Wolf, T. et al. Dynamics in protein translation sustaining T cell preparedness. Nature immunology 21, 927–937 (2020).

15 Choi, S., Ferrari, G. & Tedesco, F. S. Cellular dynamics of myogenic cell migration: molecular mechanisms and implications for skeletal muscle cell therapies. EMBO molecular medicine 12, e12357, doi:10.15252/emmm.202012357 (2020).

16 Gonzalez-Perez, O. et al. Immunological regulation of neurogenic niches in the adult brain. Neuroscience 226, 270–281, doi:10.1016/j.neuroscience.2012.08.053 (2012).

17 Suárez-Álvarez, B., López-Vázquez, A. & López-Larrea, C. Mobilization and homing of hematopoietic stem cells. Advances in experimental medicine and biology 741, 152–170, doi:10.1007/978-1-4614-2098-9_11 (2012).

18 Tanaka, T. et al. Molecular determinants controlling homeostatic recirculation and tissue-specific trafficking of lymphocytes. International archives of allergy and immunology 134, 120–134, doi:10.1159/000078497 (2004).

19 Chen, W., Dong, J., Haiech, J., Kilhoffer, M. C. & Zeniou, M. Cancer Stem Cell Quiescence and Plasticity as Major Challenges in Cancer Therapy. Stem cells international 2016, 1740936, doi:10.1155/2016/1740936 (2016).

20 Aragona, M. et al. Defining stem cell dynamics and migration during wound healing in mouse skin epidermis. Nature communications 8, 14684, doi:10.1038/ncomms14684 (2017).

21 Park, S. et al. Tissue-scale coordination of cellular behaviour promotes epidermal wound repair in live mice. Nature cell biology 19, 155–163, doi:10.1038/ncb3472 (2017).

22 Safferling, K. et al. Wound healing revised: a novel reepithelialization mechanism revealed by in vitro and in silico models. The Journal of cell biology 203, 691–709, doi:10.1083/jcb.201212020 (2013).

23 Zhao, M., Song, B., Pu, J., Forrester, J. V. & McCaig, C. D. Direct visualization of a stratified epithelium reveals that wounds heal by unified sliding of cell sheets. FASEB journal: official publication of the Federation of American Societies for Experimental Biology 17, 397–406, doi:10.1096/fj.02-0610com (2003).

24 Lang, E. et al. Coordinated collective migration and asymmetric cell division in confluent human keratinocytes without wounding. Nature communications 9, 3665, doi:10.1038/s41467-018-05578-7 (2018).

25 Angelini, T. E. et al. Glass-like dynamics of collective cell migration. Proceedings of the National Academy of Sciences of the United States of America 108, 4714–4719, doi:10.1073/pnas.1010059108 (2011).

26 Garcia, S. et al. Physics of active jamming during collective cellular motion in a monolayer. Proceedings of the National Academy of Sciences of the United States of America 112, 15314–15319, doi:10.1073/pnas.1510973112 (2015).

27 Brugués, A. et al. Forces driving epithelial wound healing. Nature physics 10, 683–690, doi:10.1038/nphys3040 (2014).

28 Möhl, C., Kirchgessner, N., Schäfer, C., Hoffmann, B. & Merkel, R. Quantitative mapping of averaged focal adhesion dynamics in migrating cells by shape normalization. Journal of cell science 125, 155–165, doi:10.1242/jcs.090746 (2012).

29 Sunyer, R. et al. Collective cell durotaxis emerges from long-range intercellular force transmission. Science (New York, N.Y.) 353, 1157–1161, doi:10.1126/science.aaf7119 (2016).

30 Huebner, R. J., Neumann, N. M. & Ewald, A. J. Mammary epithelial tubes elongate through MAPK-dependent coordination of cell migration. Development (Cambridge, England) 143, 983–993, doi:10.1242/dev.127944 (2016).

31 Alert, R. & Trepat, X. Physical models of collective cell migration. Annual Review of Condensed Matter Physics 11, 77–101 (2020).

32 Banerjee, S. & Marchetti, M. C. Continuum Models of Collective Cell Migration. Advances in experimental medicine and biology 1146, 45–66, doi:10.1007/978-3-030-17593-1_4 (2019).

33 Banerjee, S., Utuje, K. J. & Marchetti, M. C. Propagating Stress Waves During Epithelial Expansion. Physical review letters 114, 228101, doi:10.1103/PhysRevLett.114.228101 (2015).

34 Boocock, D., Hino, N., Ruzickova, N., Hirashima, T. & Hannezo, E. Theory of mechanochemical patterning and optimal migration in cell monolayers. Nature physics 17, 267–274 (2021).

35 Köpf, M. H. & Pismen, L. M. A continuum model of epithelial spreading. Soft matter 9, 3727–3734 (2013).

36 Notbohm, J. et al. Cellular Contraction and Polarization Drive Collective Cellular Motion. Biophysical journal 110, 2729–2738, doi:10.1016/j.bpj.2016.05.019 (2016).

37 Pérez-González, C. et al. Active wetting of epithelial tissues. Nature physics 15, 79–88, doi:10.1038/s41567-018-0279-5 (2019).

38 Mongera, A. et al. A fluid-to-solid jamming transition underlies vertebrate body axis elongation. Nature 561, 401–405, doi:10.1038/s41586-018-0479-2 (2018).

39 Fernández, P. A. et al. Surface-tension-induced budding drives alveologenesis in human mammary gland organoids. Nature physics 17, 1130–1136, doi:10.1038/s41567-021-01336-7 (2021).

40 Boukamp, P. et al. Normal keratinization in a spontaneously immortalized aneuploid human keratinocyte cell line. The Journal of cell biology 106, 761–771, doi:10.1083/jcb.106.3.761 (1988).

41 Serra-Picamal, X., Conte, V., Sunyer, R., Muñoz, J. J. & Trepat, X. Mapping forces and kinematics during collective cell migration. Methods Cell Biol 125, 309–330, doi:10.1016/bs.mcb.2014.11.003 (2015).

42 Preibisch, S., Saalfeld, S., Schindelin, J. & Tomancak, P. Software for bead-based registration of selective plane illumination microscopy data. Nature methods 7, 418–419, doi:10.1038/nmeth0610-418 (2010).

43 Bauer, A. et al. pyTFM: A tool for traction force and monolayer stress microscopy. PLoS computational biology 17, e1008364, doi:10.1371/journal.pcbi.1008364 (2021).

44 Weigert, M. & Schmidt, U. ZeroCostDL4Mic - What is it?, https://github.com/HenriquesLab/ZeroCostDL4Mic/wiki (2021).

45 Schmidt, U., Weigert, M., Broaddus, C. & Myers, G. Cell Detection with Star-Convex Polygons, 2018).

46 Schmidt, U., Weigert, M., Burri, O. & Haase, R. StarDist ImageJ/Fiji Plugin, https://github.com/stardist/stardist-imagej (2021).

47 Khalilgharibi, N. et al. Stress relaxation in epithelial monolayers is controlled by the actomyosin cortex. Nature physics 15, 839–847, doi:10.1038/s41567-019-0516-6 (2019).

48 Ronceray, P., Broedersz, C. P. & Lenz, M. Fiber networks amplify active stress. Proceedings of the National Academy of Sciences of the United States of America 113, 2827–2832, doi:10.1073/pnas.1514208113 (2016).

49 Wang, S. & Wolynes, P. G. Active contractility in actomyosin networks. Proceedings of the National Academy of Sciences of the United States of America 109, 6446–6451, doi:10.1073/pnas.1204205109 (2012).

50 Landau, L. D. & Lifshitz, E. M. Course of Theoretical Physics: Theory and Elasticity. Vol. 7 (Pergamon Press, 1959).

