## Supplemental information for "Mechanical coupling of supracellular stress amplification and tissue fluidization during exit from quiescence"

### Supplementary Information

#### Mathematical model

Here we give a detailed description of the model presented in the main text. We assume that the cellular monolayer is a circular layer of radius $R$ and average thickness $h_{eq}$. We further assume that the monolayer can be described as an elastic continuum with a displacement field $\mathbf{u}(\mathbf{x}, t)$ with $\mathbf{x}$ the spatial coordinate where the bold style is vector notation and $t$ is time. The displacement at a specific position is coupled to an activation concentration $c(\mathbf{x}, t)$ which regulates the contractile stresses within the cellular monolayer *e.g.* actinomyosin. Additional contributions to the model such as polarization of the cells are discussed in a following section. With negligible inertial forces and assuming no shear stress on the monolayer free surface, the in-plane force balance averaged over the monolayer height is given as

$$\mathbf{T}(\mathbf{x}, t) = h_{eq}\nabla \cdot \underline{\underline{\sigma}}(\mathbf{x}, t) \quad (1)$$

with $\mathbf{T}(\mathbf{x}, t)$ is the traction forces generated by the friction between the monolayer and the substrate and $\underline{\underline{\sigma}}(\mathbf{x}, t)$ is the cellular stress tensor. The traction force is given as $\mathbf{T}(\mathbf{x}, t) = \Gamma\partial_t\mathbf{u}(\mathbf{x}, t)$ with $\Gamma$ being the cell-substrate friction coefficient and $\partial_t$ is the time derivative operator. The cellular stress consists of the sum of the passive elastic stress and the active stress due the activation concentration $\underline{\underline{\sigma}}(\mathbf{x}, t) = \underline{\underline{\sigma_{el}}} + \underline{\underline{\sigma_a}}$. The elastic stress is modeled as a linearly elastic material $\underline{\underline{\sigma_{el}}} = E\nabla\mathbf{u}$ where $E$ is the Young's modulus. The active contribution is modeled with a linear dependence to the amount of active contractile units mimicked by the concentration field $c$ giving $\underline{\underline{\sigma_a}} = \alpha c\underline{\underline{I}}$ where $\alpha$ sets the magnitude of the contractile force and $\underline{\underline{I}}$ is the identity matrix. The dynamics of the cellular monolayer is described by the force balance between the intracellular stresses generated by the amount of activated actinomyosin ($c$) and the traction forces acting on the monolayer from the underlying substrate. Considering the monolayer to be a homogenous elastic material, the force balance in the continuum limit is expressed as

$$\Gamma\,\partial_t\mathbf{u} = h_{eq}E\nabla^2\mathbf{u} + h_{eq}\alpha\nabla c \quad (2)$$

The dynamics of the concentration field is given as a convection-reaction-diffusion equation

$$\partial_t c + (\partial_t\mathbf{u} \cdot \nabla)c = \frac{-1}{\tau_c}\left(c - c_{eq}\right) + \beta\nabla \cdot \mathbf{u} + D\nabla^2 c \quad (3)$$

$\tau_c$ is the relaxation timescale towards an equilibrium concentration $c_{eq}$ that eventually will lead to a flat homogenous stationary monolayer, $\beta$ is the rate of production of concentration due cellular contraction and $D$ is a diffusion coefficient. The terms on the left-hand side of equation (3) describes the rate change and convection of activation, respectively. On the right-hand side, the first term gives the kinetic growth/decay around an equilibrium concentration. The second term represents the increase or decay inn activation during stretching/compression of the cells and the last term describes the diffusion of the signal inside the layer.

##### *Biophysical parameters*

All model parameters are cell-type dependent and are chosen accordingly when rescaling the data obtained from the numerical simulations. The monolayer thickness, $h_{eq}$, the monolayer radius $R$ and the Young's modulus of the monolayer are measured experimentally and are listed in Supplementary Table 1. The remaining values are chosen within the same order of magnitude as reported in the literature [1,2], and their exact value are determined by comparing our numerical results with the experimentally measured time and length scales by mapping the phase space in cell layer dynamics.

**Supplementary Table 1:** Model parameters.

| Parameter | Description | Reference value | Measured value | Values in main text |
|---|---|---|---|---|
| $\Gamma$ | Cell-substrate friction coefficient | $10^7$-$10^9$ Ns m$^{-3}$ [1,2] | - | 2.5·$10^8$ Ns m$^{-3}$ |
| $E$ | Monolayer Young's modulus | - | 4000 Pa | 4000 Pa |
| $\alpha$ | Coupling coefficient between displacement and concentration | 200 Pa [2] | - | 100 Pa |
| $\tau_c$ | Concentration relaxation timescale | 4200 – 21000 s [1.2] | - | 13000 s |
| $\beta/c_{eq}$ | Production rate of concentration due cellular compression | 3·$10^{-5}$ - 6·$10^{-4}$ s$^{-1}$ [1,2] | - | 8·$10^{-5}$ s$^{-1}$ |
| $D$ | Diffusion coefficient | - | - | $10^{-9}$ m$^2$s$^{-1}$ |
| $h_{eq}$ | Average monolayer thickness at equilibrium | - | 8 µm | 8 µm |
| $R$ | Monolayer radius | - | 3.6 mm | 3.6 mm |
| $f$ | Strength of polarization-displacement coupling | 10 Pa [2] | - | 10 Pa |
| $a$ | Polarization relaxation rate | 2.16·$10^{-4}$ s$^{-1}$ [2] | - | 2.16·$10^{-4}$ s$^{-1}$ |
| $\kappa$ | Nearest neighbor alignment for **p** | 1.46·$10^{-13}$ m$^2$s$^{-1}$ [2] | - | 1.46·$10^{-13}$ m$^2$s$^{-1}$ |
| $\omega$ | Polarization alignment coefficient with the gradient of c | 3.47 ms$^{-1}$ [2] | - | 3.47 ms$^{-1}$ |

##### *Velocity scaling*

To gain some insight into the model predictions and what effect the active concentration have on the monolayer displacement we turn to a scaling analysis of equation (2). From a scaling argument we can deduce that the cellular displacement scales as

$$u\left(\frac{\Gamma}{t} - \frac{h_{eq}E}{L^2}\right) \sim \frac{h_{eq}\alpha}{L}c \qquad (4)$$

with $L$ being a characteristic length scale in the system. The velocity in the monolayer, $\sim u/t$, can then be described as

$$\frac{u}{t}\left(1-\frac{h_{eq}E}{\Gamma L^2}t\right)\sim\frac{h_{eq}\alpha}{\Gamma L}c \qquad (5)$$

At early times and small displacement we see the that the monolayer velocity scales linearly with the concentration gradient, as is recovered with a small offset in the numerical simulations and shown in Supplementary Figure 3, where the maximal velocity is plotted as a function of the initial concentration ratio $c_0$. Moreover, as the monolayer contracts the concentration in the compressed regions will be deactivated and thus reducing the monolayer velocity, and from equation (5) we see that the timescale for the velocity reduction is given by the prefactor $h_{eq}E/(\Gamma L^2)$ which for the values listed in Supplementary Table 1 yields ≈ $10^{-5}$ $s^{-1}$. This corresponds to a maximum monolayer contraction occurring at $t\approx$ 28h which is what is similar to what we observe in the experiments.

***Numerical simulations***

We non-dimensionalize equations (2)-(3) by introducing the scaling parameters $\mathbf{x}=\hat{\mathbf{x}}R$, $t=\hat{t}\Gamma R^2/(Eh_{eq})$, $\mathbf{u}=\hat{\mathbf{u}}R$ and $c=\hat{c}c_{eq}$ with hat notation indicating non-dimensional variables. The non-dimensional version of equations (2)-(3) then becomes

$$\partial_{\hat{t}}\hat{\mathbf{u}}=\hat{\nabla}^2\hat{\mathbf{u}}+\hat{\alpha}\hat{\nabla}\hat{c} \qquad (6)$$

$$\hat{\Gamma}_c\left(\partial_{\hat{t}}\hat{c}+\partial_{\hat{t}}\hat{\mathbf{u}}\cdot\hat{\nabla}\hat{c}\right)=-\hat{\tau}_c(\hat{c}-1)+\hat{\beta}\hat{\nabla}\cdot\hat{\mathbf{u}}+\hat{\nabla}^2\hat{c} \qquad (7)$$

with the dimensionless parameters $\hat{\alpha}=\alpha c_{eq}/E$ which is the ratio between the active contractile strength and the elastic stiffness, $\hat{\Gamma}_c=Eh_{eq}/(D\Gamma_u c_{eq})$ controls the strength between the elastic response and the friction due to diffusion of concentration, $\hat{\tau}_c=R^2/(D\tau_c)$ is the time scale towards an equilibrium concentration and $\hat{\beta}=\beta R^2/(Dc_{eq})$ is the ratio between the deactivation of concentration due to the monolayer compression and concentration diffusion. We solve the set of equations (6)-(7) coupled using an implicit Newton solver from the finite element library FEniCS on a circular mesh. The numerical simulations are initiated with a zero displacement condition but with a randomized concentration to mimic the stress that is generated in the monolayer during quiescence. The randomization of the initial concentration is performed such that each spatial coordinate has a 60% chance of being seeded with a value drawn from a Gaussian distribution. This initial concentration is then multiplied with a prefactor that corresponds to the starvation period in the experiments, *i.e.* large prefactor equals long starvation period. For each prefactor, yielding a ratio $\sum\hat{c}(\hat{\mathbf{x}},\ \hat{t}=0)\ /\ \sum c_{eq}=c_0$, we perform 10 simulations from which we compute the average of the system variables. The boundary conditions are set to reflect what is observed experimentally and we therefore use a homogenous Dirichlet condition on the displacement field $\hat{\mathbf{u}}(\hat{\mathbf{x}}=\partial\Omega,\ \hat{t})=0$ and a homogenous Neumann condition on the concentration, $\hat{\nabla}\hat{c}(\hat{\mathbf{x}}=\partial\Omega,\ \hat{t})\cdot\mathbf{n}=0$ with $\mathbf{n}$ being the normal vector to the domain boundary $\partial\Omega$. We multiply equations (6)-(7) with the test functions $\phi$, $\psi$ and integrate by parts over our numerical domain $\Omega$ to obtain the variational formulation of our system equations as

$$\int_{\Omega}\partial_{\hat{t}}\hat{\mathbf{u}}\cdot\phi d\hat{\mathbf{x}}+\int_{\Omega}\hat{\nabla}\hat{\mathbf{u}}\ \cdot\hat{\nabla}\phi d\hat{\mathbf{x}}-\int_{\partial\Omega}\hat{\nabla}\hat{\mathbf{u}}\cdot\mathbf{n}\phi d\mathbf{s}-\int_{\Omega}\hat{\alpha}\hat{\nabla}\hat{c}\cdot\phi d\hat{\mathbf{x}}=0, \quad (8)$$

$$\int_{\Omega}\hat{\Gamma}_c\big(\partial_{\hat{t}}\hat{c}+\partial_{\hat{t}}\hat{\mathbf{u}}\cdot\hat{\nabla}\hat{c}\big)\psi d\hat{\mathbf{x}}+\int_{\Omega}\hat{\tau}_c(\hat{c}-1)\psi d\hat{\mathbf{x}}-\int_{\Omega}\hat{\beta}\hat{\nabla}\cdot\ \hat{\mathbf{u}}\psi d\hat{\mathbf{x}}+\int_{\Omega}\hat{\nabla}\hat{c}\cdot\hat{\nabla}\psi d\hat{\mathbf{x}}=0. \quad (9)$$

As our equations are reduced to first order equations, we discretize them using linear elements with the implicit discrete time derivatives as

$$\frac{\hat{\mathbf{u}}^n-\hat{\mathbf{u}}^{n-1}}{\Delta\hat{t}}=f(\hat{\mathbf{u}}^n,\hat{c}^n) \quad (10)$$

$$\frac{\hat{c}^n-\hat{c}^{n-1}}{\Delta\hat{t}}=g(\hat{\mathbf{u}}^n,\hat{c}^n) \quad (11)$$

with $n$ being the current time step we evaluate, $\Delta\hat{t}$ is the time step spacing and $f, g$ represents the right-hand side of equations (8)-(9), respectively.

**Supplementary Table 2*:*** Non-dimensional model parameters.

| Parameter | Description | Tested value range | Values in main text |
|---|---|---|---|
| $\hat{\alpha}$ | Ratio between contractile strength and elastic stiffness | 0.01 - 10 | 10 |
| $\hat{\beta}$ | Deactivation of concentration due to monolayer compression | $10^{-2}$ - $10^{4}$ | 1 |
| $\hat{\Gamma}_c$ | Strength between elastic response and substrate friction | $10^{-4}$ - $10^{2}$ | 100 |
| $\hat{\tau}_c$ | Concentration relaxation timescale | $10^{-2}$ - $10^{2}$ | 1 |
| $\hat{f}$ | Displacement-polarization coupling coefficient | 1.125 | 1.125 |
| $\hat{a}$ | Polarization relaxation timescale | 22 | 2 |
| $\hat{\kappa}$ | Strength of nearest neighbor alignment | $10^{-3}$ | $10^{-3}$ |
| $\hat{\omega}$ | Polarization alignment rate with concentration gradient | 0.975 | 0.975 |

***Parameter sensitivity - contraction center formation***

To determine the effect from the non-dimensional system parameters on the dynamics we performed a parameter sensitivity study. All simulations are initialized with the same initial condition depicted in Supplementary Figure 5 with a ratio $c_0 = 1.2$.

The results from the study is shown in Supplementary Figures 6, 7, 8, with the red dotted lines highlighting the parameter range that yields collective motion and formation of a contraction center, and is summarized in the following points:

- $\hat{\alpha}$ determines the magnitude of the monolayer displacement for a given value of $c_0$. A sufficiently large value of $\hat{\alpha}$ is crucial to obtain the large scale migration needed to form one big central contraction center.

• The dynamics is unaffected by small values of $\hat{\beta}$ yet it can be important with large values of $\hat{\alpha}$ to control the maximum central compression. For large values of $\hat{\beta}$ the concentration deactivation due to the monolayer compression can stop the collective migration and prevent the formation of a central contraction center.

• Smaller values of $\hat{\Gamma}_c$ lead to smoother and more symmetric migration due to diffusion of the active contractile units yet with a severely reduced displacement field magnitude. Larger values of $\hat{\Gamma}_c$ increases the displacement magnitude but not to the same extent as $\hat{\alpha}$. A sufficiently large value of $\hat{\Gamma}_c$ is needed to obtain the convective dynamics observed in the experiments.

• Large values of $\hat{\tau}_c$ increase the deactivation rate of the concentration to such an extent that collective migration can be halted but very large values are needed to significantly affect the dynamics.

To extend our analysis we plot the maximum monolayer velocity magnitude, $\max(|\partial_{\hat{t}}\hat{\mathbf{u}}|)$, from the simulation data that provided the displacement fields in Supplementary Figures 6-8, shown in Supplementary Figure 9. The previously summarized conclusions are further enhanced and we see that the velocity magnitude is approximately unaffected by changes in both $\hat{\tau}_c$ and $\hat{\beta}$. Moreover, we see that the monolayer velocity magnitude increases linearly with $\hat{\alpha}$ and as the square root of $\hat{\Gamma}_c$, making them both important to achieve large scale collective migration.
We can thus separate the four dimensionless parameters into two categories, with $\hat{\alpha}$ and $\hat{\Gamma}_c$ being the parameters important for initiating the collective migration and $\hat{\tau}_c$ and $\hat{\beta}$ are important to slow down the migration at late times, preventing possible diverging compression in the monolayer, and to enforce the inevitable relaxation to a flat homogenous equilibrium monolayer at $\hat{t} \rightarrow \infty$.

***Numerical solution in square geometry***
In order to check if the collective migration towards the center of the circular mesh is a feature induced by the geometry, we solve the same system on a square mesh. In Supplementary Figure 10a the displacement field in the square geometry shows that the formation of a large central contraction center occurs also for non-circular meshes. Moreover, the displacement field magnitude only differs by $\sim$ 10% to that in the circular mesh (Supplementary Figure 10b) given the same non-dimensional parameters $\hat{\alpha} = \hat{\beta} = \hat{\tau}_c = \hat{\Gamma}_c = 1$.

***Model and experiments - comparison***
We performed numerical simulations using a wide range of initial concentration ratios and verified that an initial concentration can cause a collective cellular migration towards the center of the monolayer. The initial response to the random concentration is to form small local contraction centers of high cell density where the peaks in the initial concentration are located, accompanied by a large traction force pointing in towards these centers. After these initial

density center forms, the monolayer displacement is directed inwards toward the center of the monolayer where one large cell density center is formed. Below a threshold ratio value $c_0$, there is no collective motion towards the center but the monolayer relaxes to equilibrium after the initial formation of the small density centers (Supplementary Video 6). Around the threshold ratio value there are stronger local density centers that are formed with a subsequent collective motion towards the center of the mesh. However, with small gradients in the concentration the dynamics stagnates before a single central contraction center is formed (Supplementary Video 8). Above the threshold value there is a grand collective migration towards the center of the mesh after the initial formation of small local contraction centers. Now the dynamics does not stagnate until a high density central contraction center is formed (Supplementary Video 7). The dynamics are recovered in the experiments (Supplementary Video 2) and there are two key parameters we can extract. The traction force is shown to peak at the early formation of the small local contraction centers and decay as the cells collective migration begins. The results from the numerical simulations are shown in Figure 4g that also highlights that the traction force magnitude increase with the initial concentration ratio $c_0$. The overall traction force profile is qualitatively similar to that measured in the experiments, shown in Figure 2f-g. The increased traction force magnitude also indicates an increased migration velocity. The averaged radial migration velocity calculated from the numerical data is shown in Figure 4h and is in good agreement with the experimental results shown in Figure 4j.

***Polarization effects***

From the experiments there is no indication that the collective motion and formation of a contraction center is driven by cellular polarization. However, we know that cellular polarization can have an effect on the monolayer dynamics [1-7].

To investigate the effects that cellular polarization has on the dynamics we introduce a polarization vector $\mathbf{p}(\mathbf{x}, t)$. Following [2], the dynamics of the polarization field can be described by the following equation

$$\partial_t \mathbf{p} = a(1 - |\mathbf{p}^2|)\mathbf{p} + \kappa \nabla^2 \mathbf{p} + \omega \nabla c \quad (12)$$

with $a$ controls the rate of relaxation towards a homogeneously polarized monolayer, $\kappa$ is the strength of the nearest neighbor alignment and $\omega$ is the coupling coefficient to the concentration gradient such that local cell polarization points towards regions of high concentration. Further, we couple the polarization field to the monolayer displacement through the traction force which now reads $\mathbf{T}(\mathbf{x}, t) = \Gamma \partial_t \mathbf{u}(\mathbf{x}, t) - f\mathbf{p}(\mathbf{x}, t)$, with $f$ being the coupling coefficient between the cell polarization and the monolayer displacement. Inserted into equation (1) we get

$$\Gamma \partial_t \mathbf{u} = h_{eq} E \nabla^2 \mathbf{u} + h_{eq} \alpha \nabla c + f\mathbf{p} \quad (13)$$

We non-dimensionalize equations (12)-(13) as in the Numerical methods section to obtain

$$\partial_{\hat{t}} \hat{\mathbf{u}} = \hat{\nabla}^2 \hat{\mathbf{u}} + \hat{\alpha} \hat{\nabla} \hat{c} + \hat{f} \hat{\mathbf{p}} \quad (14)$$

and

$$\partial_{\hat{t}}\hat{\mathbf{p}} = \hat{\alpha}(1 - |\hat{\mathbf{p}}|^2)\hat{\mathbf{p}} + \hat{\kappa}\hat{\nabla}^2\hat{\mathbf{p}} + \hat{\omega}\hat{\nabla}\hat{c}. \quad (15)$$

with $\hat{f} = fR/(h_{eq}E)$ being the strength of the polarization-displacement coupling, $\hat{a} = a\tau$ the polarization relaxation rate, $\hat{\kappa} = \kappa\tau/R^2$ the local neighbor alignment coefficient and $\hat{\omega} = \omega\tau c_{eq}/R$ is the strength of the polarization alignment with the concentration gradient. We now have a non-dimensional coupled three equation system consisting of equations (14), (7), (15) that we solve as described in the Numerical methods section. Using parameter values found in literature, see Supplementary Table 1 (Supplementary Table 2 for non-dimensional units), we see the results from a numerical simulation in Supplementary Figure 11. At three different times we compare the displacement field with a polarization field to the displacement field without a polarization field using the same initial condition displayed in Supplementary Figure 5. There are no significant differences in displacement magnitude, length scale or time scale between polarized and unpolarized results. This indicate that cellular polarization have no significant impact on the large scale collective migration observed in this study which is consistent with the experiments and we use this as a justification to neglect this effect in our theoretical model.

#### Supplementary Figures

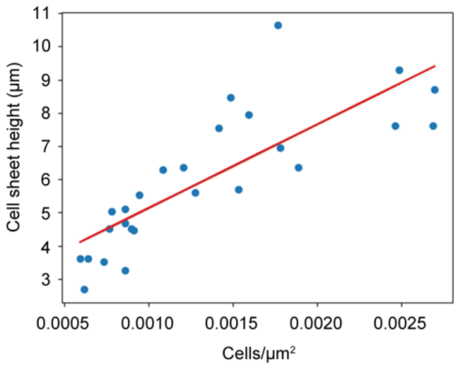


**Supplementary Figure 1:** Monolayer thickness measurements. Manual measurements of cell sheet thickness were performed on confluent keratinocyte monolayers subjected to serum depletion for two days followed by serum re-stimulation. Graph shows average values (blue dots) of cell sheet thickness at different cell densities. The red line shows the best fit curve based on linear regression.

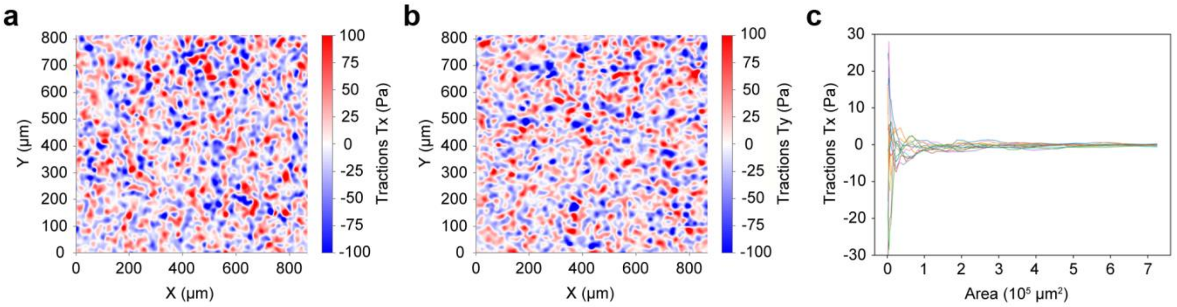


**Supplementary Figure 2:** Random orientation of local traction forces. **a**,**b** Plotting of individual traction force components $T_x$ (**a**) and $T_y$ (**b**) across a microscopic field of view (866x814 µm). **c**, Plot showing the average of the traction force component $T_x$ within a progressively increasing microscopic field area. The time point selected is 1 h after serum stimulation.

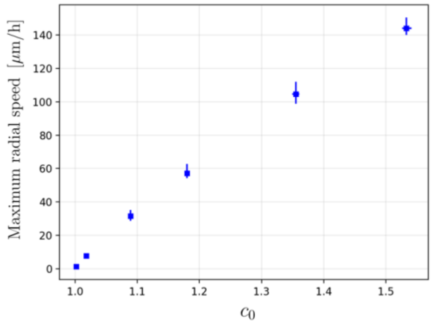


**Supplementary Figure 3:** Maximum radial velocity in the monolayer as calculated from the numerical simulations. The maximum velocity follows a linear trend with the initial concentration magnitude with a small decaying offset.

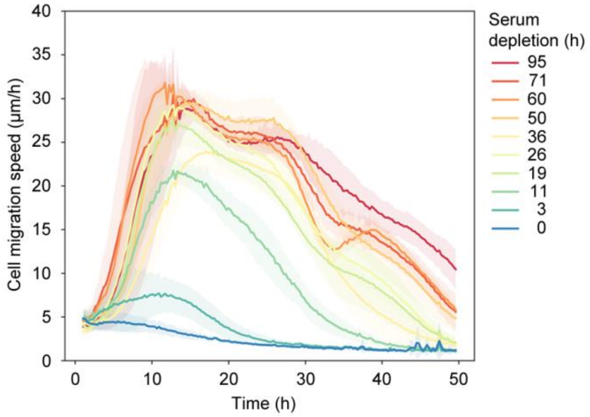


**Supplementary Figure 4:** Line plots showing changes in cell sheet migration speed over time after different time periods of serum depletion prior to serum re-activation of confluent cell sheets. Graph shows mean values ± SD.

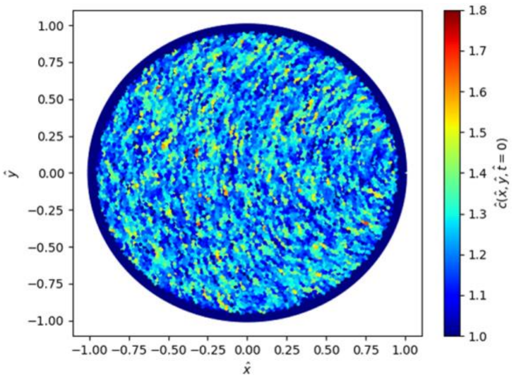


**Supplementary Figure 5:** Initial condition for the concentration $\hat{c}(\hat{\mathbf{x}}, \hat{t} = 0)$ for all the simulations in the parameter study. The initial condition has a ratio $c_0$ = 1.2.

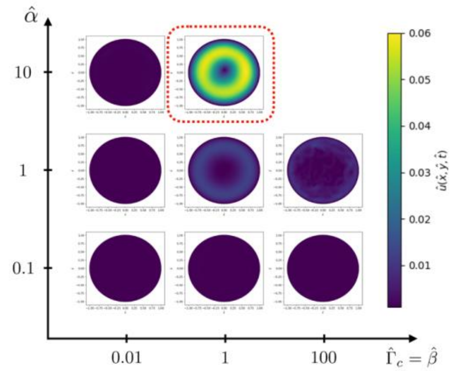


**Supplementary Figure 6:** Displacement field $\hat{\mathbf{u}}(\hat{\mathbf{x}}, \hat{t})$ at the time of maximal contraction plotted as a function of $\hat{\Gamma}_c$ and $\hat{\tau}_c$. In all the simulations $\hat{\tau}_c = 1$. The red dashed line marks the parameter space for which a global contraction center is formed.

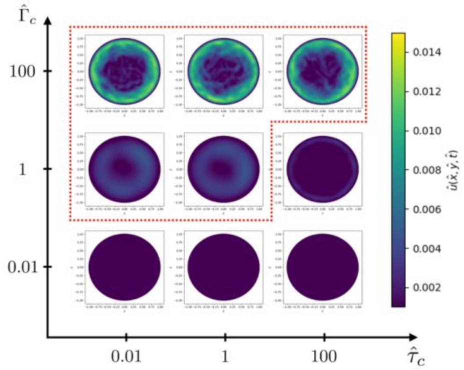


**Supplementary Figure 7:** Displacement field $\hat{\mathbf{u}}(\hat{\mathbf{x}}, \hat{t})$ at the time of maximal contraction plotted as a function of $\hat{\Gamma}_c$ and $\hat{\tau}_c$. In all the simulations $\hat{\beta} = 0$ and $\hat{\alpha} = 1$. The red dashed line marks the parameter space for which a global contraction center is formed.

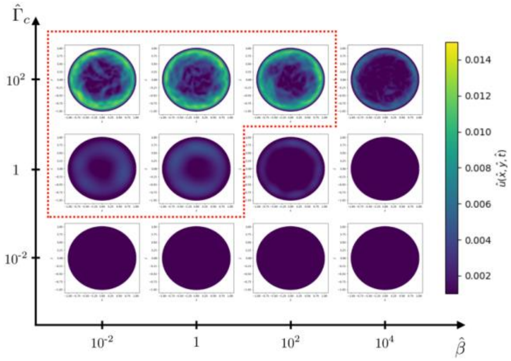

**Supplementary Figure 8:** Displacement field $\hat{\mathbf{u}}(\hat{\mathbf{x}}, \hat{t})$ at the time of maximal contraction plotted as a function of $\hat{\Gamma}_c$ and $\hat{\beta}$. In all the simulations $\hat{\alpha} = 1$ and $\hat{\tau}_c = 0$. The red dashed line marks the parameter space for which a global contraction center is formed.

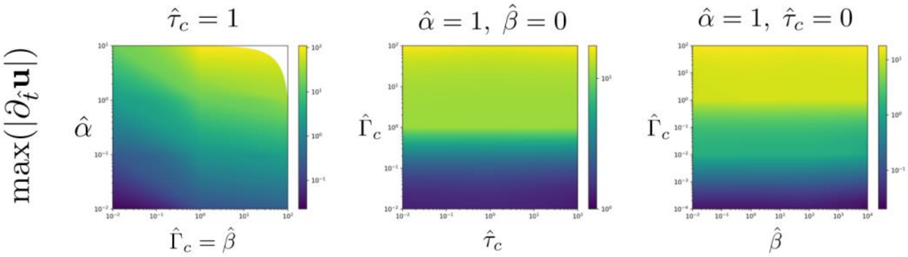

**Supplementary Figure 9:** Maximum value of the monolayer velocity magnitude, $\max(|\partial_{\hat{t}}\hat{\mathbf{u}}|)$, corresponding to parameter study simulations depicted in Supplementary Figures 6-8.

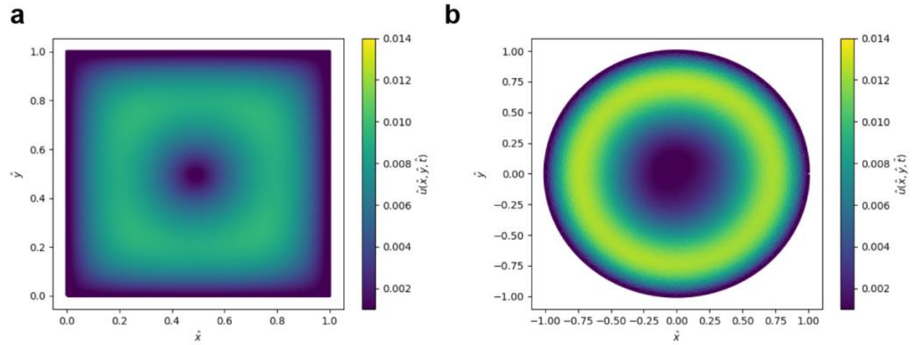

**Supplementary Figure 10:** Supplementary Figure 7: Displacement field $\hat{\mathbf{u}}(\hat{\mathbf{x}}, \hat{t})$ with $\hat{\alpha} = \hat{\beta} = \hat{\tau}_c = \hat{\Gamma}_c = 1$ in a square mesh (**a**) and a circular mesh (**b**).

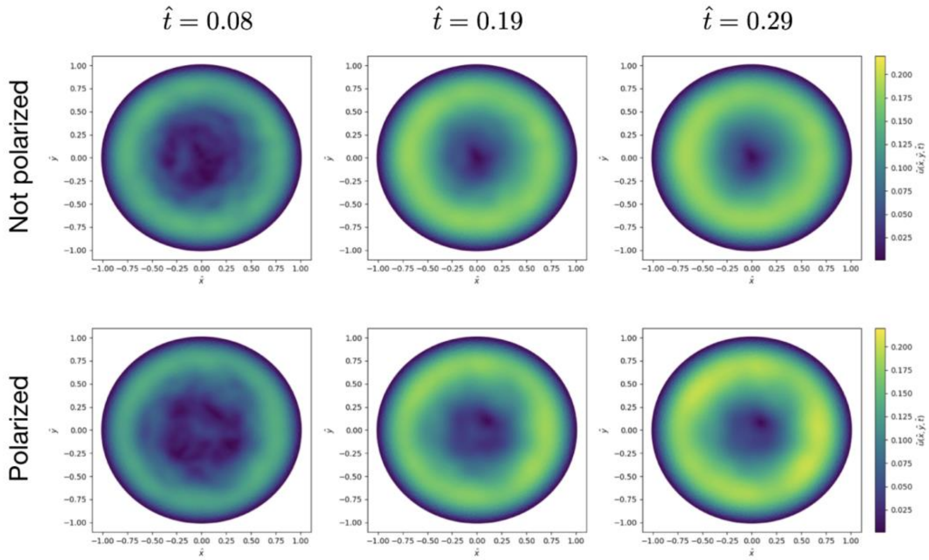


**Supplementary Figure 11:** Displacement field at three different times from a simulation with an unpolarized monolayer, i.e. $\hat{f}= 0$ (top row), and with $\hat{f}= 1.125$ (bottom row).

#### Supplementary Videos

##### Supplementary Video 1

Movies showing cell sheet dynamics in 96-well glass bottom plates. Left panel: proliferation (HaCaT keratinocytes without treatment before imaging); Middle panel: quiescence (HaCaT keratinocytes serum-depleted for 48 h before imaging); Right panel: fluidization (HaCaT keratinocytes serum-depleted for 48 h and subsequently serum re-stimulated before imaging). Microscopy settings: 4x air objective, binning 2 (3.367x3.367 µm pixel size), a time interval of 16 min and an imaging period time of 30 h. The movie covers an imaging period of 27 h and every second frame is shown. Each frame is composed of 4 tailed images. Scale bar, 1 mm.

##### Supplementary Video 2

Visualization of serum-stimulated cell sheet dynamics based on cell flow (left panel), cell density (middle panel), and cell sheet thickness (right panel). Microscopy settings: 4x air objective, binning 2 (3.367x3.367 µm pixel size), a time interval of 16 min and an imaging period time of 30 h. The movie covers an imaging period of 27 h and every second frame is shown. Each frame is composed of 4 tailed images.

##### Supplementary Video 3

Basal actin dynamics 0 to 2 h after serum stimulation. HaCaT cells expressing LifeAct-RFP were subjected to serum deprivation for 48 h and subsequently re-stimulated with serum. A single confocal z-plane representing the basal cell surface is shown. The movie shows the two first hours after serum stimulation. The frame interval is set to 2 min between frames.

##### Supplementary Video 4

Basal actin dynamics 15 to 16 h after serum stimulation. HaCaT cells expressing LifeAct-RFP were subjected to serum deprivation for 48 h. Cells were then re-stimulated with serum for 15 h before imaging. A single confocal z-plane representing the basal cell surface is shown. The movie shows basal actin dynamics between 15 and 16 h after serum re-stimulation. The frame interval is set to 1 min between frames.

**Supplementary Video 5**
Traction force time lapse microscopy of a serum-stimulated confluent quiescent cell sheet. Phase contrast (left panel), traction forces (middle panel), and intercellular tension (right panel) are depicted. Microscopy settings: 10x FLUAR objective (1331x1331 µm field of view), binning 1. A time interval of 16 min and a total imaging period of 18 h are shown.

**Supplementary Video 6**
Time lapse of the cellular density in the monolayer obtained from a numerical simulation using a normalized initial concentration $c(\mathbf{x}, t = 0)/c_{eq} = 0.1$. We observe that there is a small initial rearrangement in the monolayer density due to the inhomogeneous concentration. Due to the small gradients in the concentration there is no collective motion and the monolayer quickly adopts a quasi-static homogenous profile.

**Supplementary Video 7**
Time lapse of the cellular density in the monolayer obtained from a numerical simulation using a normalized initial concentration $c(\mathbf{x}, t = 0)/c_{eq} = 1.0$. After the formation of many strong local contraction centers there is a collective cell migration response towards the center of the monolayer. This collective migration results in the formation of one large global contraction center.

**Supplementary Video 8**
Time lapse of the cellular density in the monolayer obtained from a numerical simulation using a normalized initial concentration $c(\mathbf{x}, t = 0)/c_{eq} = 0.5$. At higher concentration levels we observe coordinated cellular motion in the monolayer after the initial formation of small local contraction centers. However, the concentration gradients are not large enough to stimulate the formation of a global contraction center.

**References**
1 Banerjee, S., Utuje, K. J. & Marchetti, M. C. Propagating Stress Waves During Epithelial Expansion. *Physical review letters* **114**, 228101, doi:10.1103/PhysRevLett.114.228101 (2015).
2 Notbohm, J. *et al.* Cellular Contraction and Polarization Drive Collective Cellular Motion. *Biophysical journal* **110**, 2729-2738, doi:10.1016/j.bpj.2016.05.019 (2016).
3 Bi, D., Lopez, J., Schwarz, J. M. & Manning, M. L. A density-independent rigidity transition in biological tissues. *Nature Physics* **11**, 1074-1079 (2015).
4 Boocock, D., Hino, N., Ruzickova, N., Hirashima, T. & Hannezo, E. Theory of mechanochemical patterning and optimal migration in cell monolayers. *Nature Physics* **17**, 267-274 (2021).
5 Köpf, M. H. & Pismen, L. M. A continuum model of epithelial spreading. *Soft matter* **9**, 3727-3734 (2013).
6 Loewe, B., Serafin, F., Shankar, S., Bowick, M. J. & Marchetti, M. C. Shape and size changes of

adherent elastic epithelia. *Soft Matter* **16**, 5282-5293, doi:10.1039/d0sm00239a (2020).
7 Pérez-González, C. *et al.* Active wetting of epithelial tissues. *Nat Phys* **15**, 79-88, doi:10.1038/s41567-018-0279-5 (2019).
